# A non-canonical CDK, Pho85 regulates the restart of the cell-cycle following stress

**DOI:** 10.1101/2024.08.27.609989

**Authors:** Natsuko Jin, Yui Jin, Yu Oikawa, Akihiko Nakano, Yoshinori Ohsumi, Lois S. Weisman

**Author notes:** Lead Contact: Lois Weisman.

## Abstract

Environmental stress induces an arrest of the cell cycle. Thus, release from this arrest is essential for cell survival. The cell-cycle-arrest occurs via the down regulation of the cyclins that drive the main cyclin dependent kinase, CDK1/Cdc28. However, it was not clear how cells escape this potentially fatal arrest. Here we show that prior to the restoration of CDK1/Cdc28 cyclins, a non-canonical CDK, Pho85, initiates a cascade to restart the cell cycle. We demonstrate that following stress, Pho85 phosphorylates the Sch9 kinase, which in turn directly phosphorylates the transcriptional inhibitor Whi5, the yeast analog of RB1/retinoblastoma, and a CDK1 target. This promotes Whi5 translocation from the nucleus, and the release of the stress-induced arrest at G_1_ phase. In addition, we find that in parallel with Pho85, CDK1/Cdc28 also plays a role in the control of Whi5. Together, these findings provide insights into how cells re-enter the cell cycle during recovery from stress and reveal that a non-canonical CDK and cyclin takes on essential roles and acts via a pathway that functions in parallel with CDK1/Cdc28.

## Introduction

The precise regulation of the cell-cycle is essential for cell proliferation and viability and is fundamental for proper development and tissue maintenance in all organisms. In eukaryotic cells, the cell-cycle is regulated by cyclin-dependent kinase (CDK)-cyclin complexes (Rhind and Russell, 2012). In budding yeast, a single CDK, Cdc28, the yeast homolog of mammalian CDK1, regulates the cell-cycle (Nasmyth, 1993). Cdc28 activity is tightly controlled via the transient expression of cyclins at specific phases of the cell-cycle. The precise timing of cyclin expression coordinates progression through each cell-cycle phase (Bloom and Cross, 2007). The earliest event is the commitment to undergo cell division. This occurs in early G_1_ phase, and in yeast is also referred to as START. One key player in START is Whi5, which is initially hypo-phosphorylated by both CDK- and non-CDK kinases at early G_1_ phase. These phospho-sites are required for further CDK dependent hyper-phosphorylation at late G_1_ phase, which drives the export of Whi5 from the nucleus (Xiao et al., 2024). It was initially thought that this latter phosphorylation of Whi5 was carried out by Cdc28-Cln3 (Costanzo et al., 2004; de Bruin et al., 2004), however, more recently it was found that that Cdc28-Cln1 and Cdc28-Cln2 are responsible for Whi5 hyperphosphorylation in late G_1_ phase (Kõivomägi et al., 2021). Importantly, phosphorylation of Whi5 and its translocation from the nucleus is a robust indicator for progression from early G_1_ phase (Kõivomägi et al., 2021).

Cdc28-Cln3 also has roles in progression through START via its phosphorylation of RNA polymerase (Kõivomägi et al., 2021), and via Cln3 regulation of the levels of Cln1 and Cln2 in a post-transcriptional mechanism that is independent of Whi5 (Brambila et al., 2024).

Whi5 is similar in function to the mammalian tumor suppressor, RB1/retinoblastoma (RB transcriptional corepressor 1). Whi5 cycles between the nucleus and cytoplasm, and is a negative transcriptional regulator of genes required for the G_1_-S phase transition, including the Cdc28 cyclins, Clb5 and Clb6 (Costanzo et al., 2004; de Bruin et al., 2004). Whi5 export releases the negative transcriptional regulation of these genes (Costanzo et al., 2004; de Bruin et al., 2004). In addition, cell growth-dependent dilution of Whi5 may also play a critical role in progression of the cell-cycle (Palumbo et al., 2016; Schmoller et al., 2015; Talarek et al., 2017), however, this remains an ongoing area of investigation (Schmoller et al., 2015; Sommer et al., 2021).

The cell-cycle machinery is also regulated by environmental cues. In eukaryotes, environmental stresses including hyperosmotic stress, nutrient limitation and high temperature cause an arrest of the cell-cycle at multiple cell-cycle phases (Argüello-Miranda et al., 2022; Barbet et al., 1996; Clotet et al., 2006; Escoté et al., 2004; Joaquin et al., 2012; Johnston and Singer, 1980; Matsui et al., 2013). At G_1_ phase, stress induced-arrest of the cell-cycle was proposed to be due to inactivation of Cdc28 via transcriptional downregulation of G_1_-specific cyclins (Belli et al., 2001; Escote et al., 2004). Importantly, this stress-induced cell-cycle arrest at G_1_ phase is transient. These findings strongly suggest that following stress, additional pathways may be critical to restart the cell-cycle. However, the mechanisms to restart the cell-cycle following stress were unknown.

Here we discover that a non-canonical pathway, which is activated by the CDK5/Pho85, restarts the cell-cycle specifically at the G_1_-S transition. Pho85 initiates a cascade via direct phosphorylation and positive regulation of an AGC kinase, Sch9, which in the canonical cell-cycle is positively regulated via phosphorylation by TORC1 (Urban et al., 2007a). We show that Sch9 then directly acts to phosphorylate Whi5, which results in the export of Whi5 from the nucleus. This is required for the restart of the cell cycle at G_1_ phase and for progression through the G_1_/S transition. We show that this non-canonical pathway is triggered by two distinct stresses, hyperosmotic shock and recovery from nutrient starvation. These findings predict that the non-canonical CDK, Pho85 pathway is required for cellular homeostasis under multiple stress conditions, through the regulation of cell-cycle progression at G_1_ phase.

## Results

### Stress induces a transient arrest at multiple phases of the cell-cycle

At G_1_ phase, hyperosmotic stress induces an arrest of the cell-cycle due to inactivation of Cdc28, which occurs via transcriptional downregulation of the G_1_-specific cyclins, Cln1, Cln2 and Cln3 (Bellí et al., 2001; Escoté et al., 2004). We tested and found that the levels of cyclins which function at other phases of the cell-cycle, were also decreased following stress. We treated wild-type cells with hyperosmotic stress at different points of the cell-cycle and assessed protein levels of cyclins for Cdc28. We found that in addition to the G_1_-specific cyclins Cln1, Cln2 and Cln3, the expression of G_2_-M phase-specific cyclins Clb1 and Clb2 and the G_1_-S transition cyclins Clb5 and Clb6 were inhibited (Fig 1A-E). The fact that the levels of all cyclins tested were rapidly lowered following stress indicates that Cdc28 is inactivated at multiple phases of the cell-cycle.

**Figure 1.**
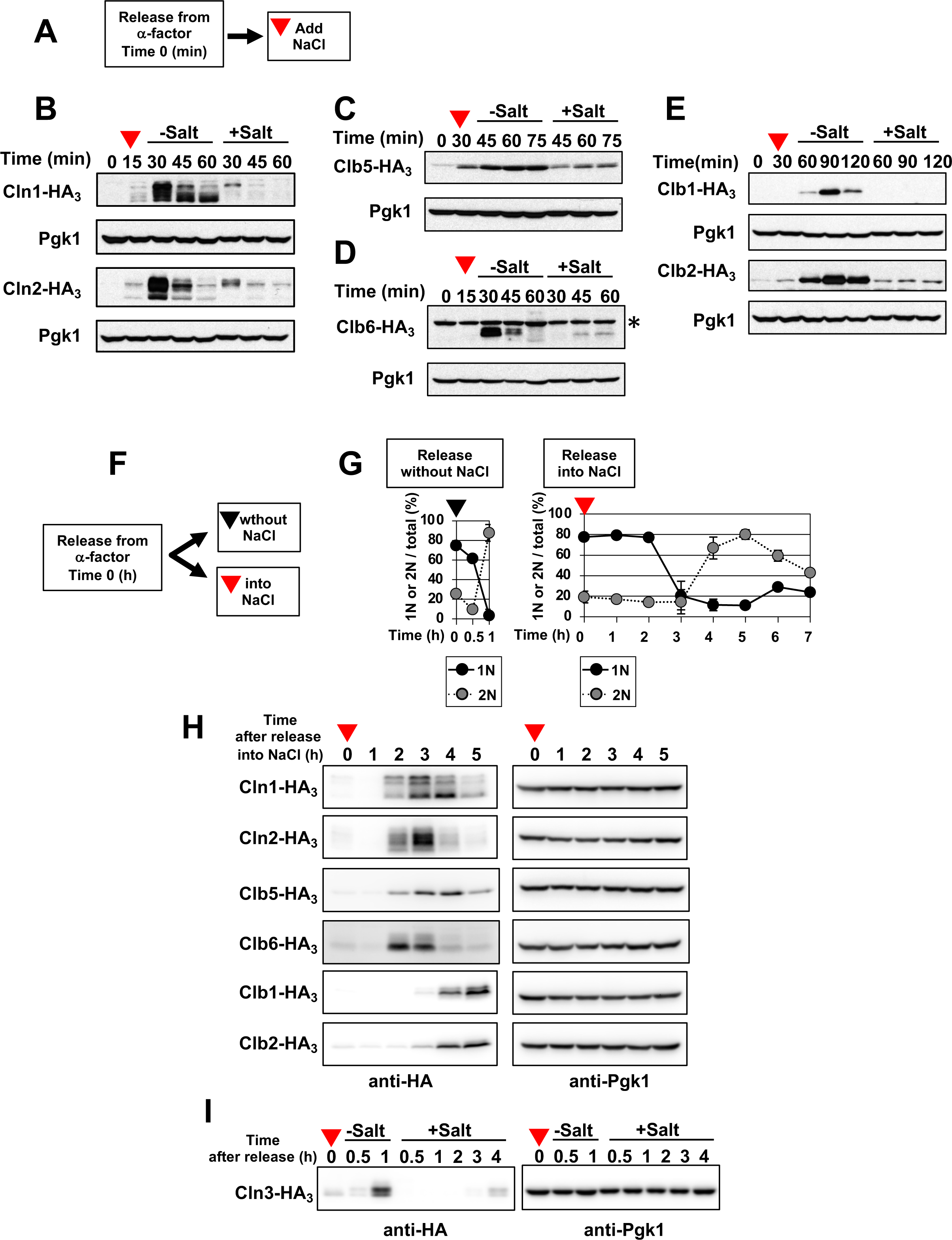
Hyperosmotic stress causes a transient arrest of cell-cycle progression. (A) Schematic of experimental design for (B)-(E). (B)-(E) The levels of each CDK1/Cdc28 cyclin tested were greatly diminished following hyperosmotic stress, which suggests that Cdc28 is not active under these conditions. Yeast expressing Cln1-HA_3_, Cln2-HA_3_, Clb1-HA_3_, Clb2-HA_3_, Clb5-HA_3_ or Clb6-HA_3_ were arrested with α-factor (Zymo Research) with 80% of the population with 1N DNA content, and were released to fresh medium, then treated with high salt medium (0.9 M NaCl) at the times indicated. Red triangle, time point of adding salt to the medium. Black asterisk in (D) indicates background band. Western blot: anti-HA or anti-Pgk1 (control). Blots representative of 3 independent experiments. (F) Schematic of experiment for (G)-(I). (G) Wild-type yeast arrested with α-factor with 80% of the population with 1N DNA content, were released into fresh medium with or without 0.9 M NaCl. DNA content was measured by flow cytometry at the times indicated. Mean values ± SD. (H) and (I) The levels of each CDK1/Cdc28 cyclin tested were transiently downregulated following hyperosmotic stress. Concomitant of the restart of cell-cycle, Cdc28 cyclins were re-expressed in sequence. Yeast expressing Cln1-HA_3_, Cln2-HA_3_, Cln3-HA_3_, Clb1-HA_3_, Clb2-HA_3_, Clb5-HA_3_, or Clb6-HA_3_ were arrested with α-factor with 80% of the population with 1N DNA content, and were released to high salt medium (0.9 M NaCl). Blots representative of 3 independent experiments.

Notably, the stress-induced cell-cycle arrest is transient. We observed that following hyperosmotic stress, wild-type cells can exit from the stress-induced arrest from at least two phases of the cell-cycle, G_1_ phase and G_2_-M phase. When haploid yeast cells are treated with α-factor, they arrest in early G_1_ phase (START) and have 1N DNA content. When cells exit START and progress through S phase and complete DNA synthesis, the cells have 2N content (G_2_-M phase). To test the ability of cells to recover from an arrest in G_1_ phase, we arrested cells with α-factor and then released the cells into fresh medium with or without high salt (0.9 M NaCl) (Fig 1F and 1G). At 1 h after release into normal medium, cells progressed through G_1_ phase, completed DNA synthesis and entered into G_2_ and M phases of the cell-cycle (i.e., had predominantly 2N DNA content; Fig 1G). In contrast, following 2 h in hyperosmotic stress, ∼80% of the cells continued to have 1N DNA content, indicating that the cells were arrested at G_1_ phase (Fig 1G) (Bellí et al., 2001; Clotet et al., 2006; Escoté et al., 2004). However, after 3 h in hyperosmotic stress, cells exited the arrest and progressed to G_2_ and M phases (Fig 1G). In addition to recovery from an arrest in G_1_ phase during hyperosmotic stress, G_1_-specific cyclins Cln1and Cln2, the G_1_-S transition cyclins Clb5 and Clb6, and G_2_-M phase-specific cyclins Clb1 and Clb2 were re-expressed in a sequential manner (Fig 1H). Next, to test the ability of cells to recover from an arrest in G_2_ or M phase, we treated wild-type cells with α-factor, released cells to fresh medium and incubated for an additional 1 h until ∼90% of cells had 2N DNA content (G_2_ or M phase). Then cells were exposed to hyperosmotic stress (Supplemental Fig 1A and 1B). Following 2 h in hyperosmotic stress, cells continued to have 2N DNA content, indicating that the cells remained arrested at G_2_ and M phases (Supplemental Fig 1B). Moreover, the G_2_-M phase-specific cyclins Clb1 and Clb2 were expressed at the one and two hour timepoints (Supplemental Fig 1C). Eventually, following 4 h in hyperosmotic stress, the G_1_-specific cyclins Cln1, Cln2 and Cln3 were re-expressed (Supplemental Fig 1C) and after 5 h, many of the cells completed mitosis as assessed by 1N DNA content (Supplemental Fig 1B). Taken together, these observations suggest that hyperosmotic stress causes the transient arrest of the cell-cycle at multiple phases.

### A non-canonical CDK, Pho85 is required to restart the cell-cycle in G_1_ phase following hyperosmotic stress

CDK1/Cdc28-Cln3 drives START by phosphorylating the RNA polymerase II subunit Rbp1 to promote entry to the cell-cycle (Kõivomägi et al., 2021). Notably, we found that following 2 h in hyperosmotic stress, Cln3 remains absent even after the cell-cycle resumes (Fig 1I). However, despite the absence of Cln3, the re-expression of selected CDK1/Cdc28 cyclins occurs, specifically Clb5, and Clb6, G_1_-S specific cyclins (Fig 1H). These findings suggest that following stress, instead of Cdc28-Cln3, other mechanisms restart the cell-cycle.

Based on previous studies, we considered the non-canonical CDK, Pho85, as a potential candidate to drive the restart of the cell-cycle during stress. Pho85 functions with ten cyclins and plays roles in signaling during several types of stress (Huang et al., 2007). Pho85 is required for cell-cycle progression during phosphate limitation (Menoyo et al., 2013) and for growth of yeast under hyperosmotic stress (Jin et al., 2017) (Fig 2A). However, under basal conditions, the contributions of Pho85 (Fig 2B) and some of its cyclins including Pcl1, Pcl2, Pcl7 and Pcl9 are not essential (Bállega et al., 2019; Huang et al., 2009).

**Figure 2.**
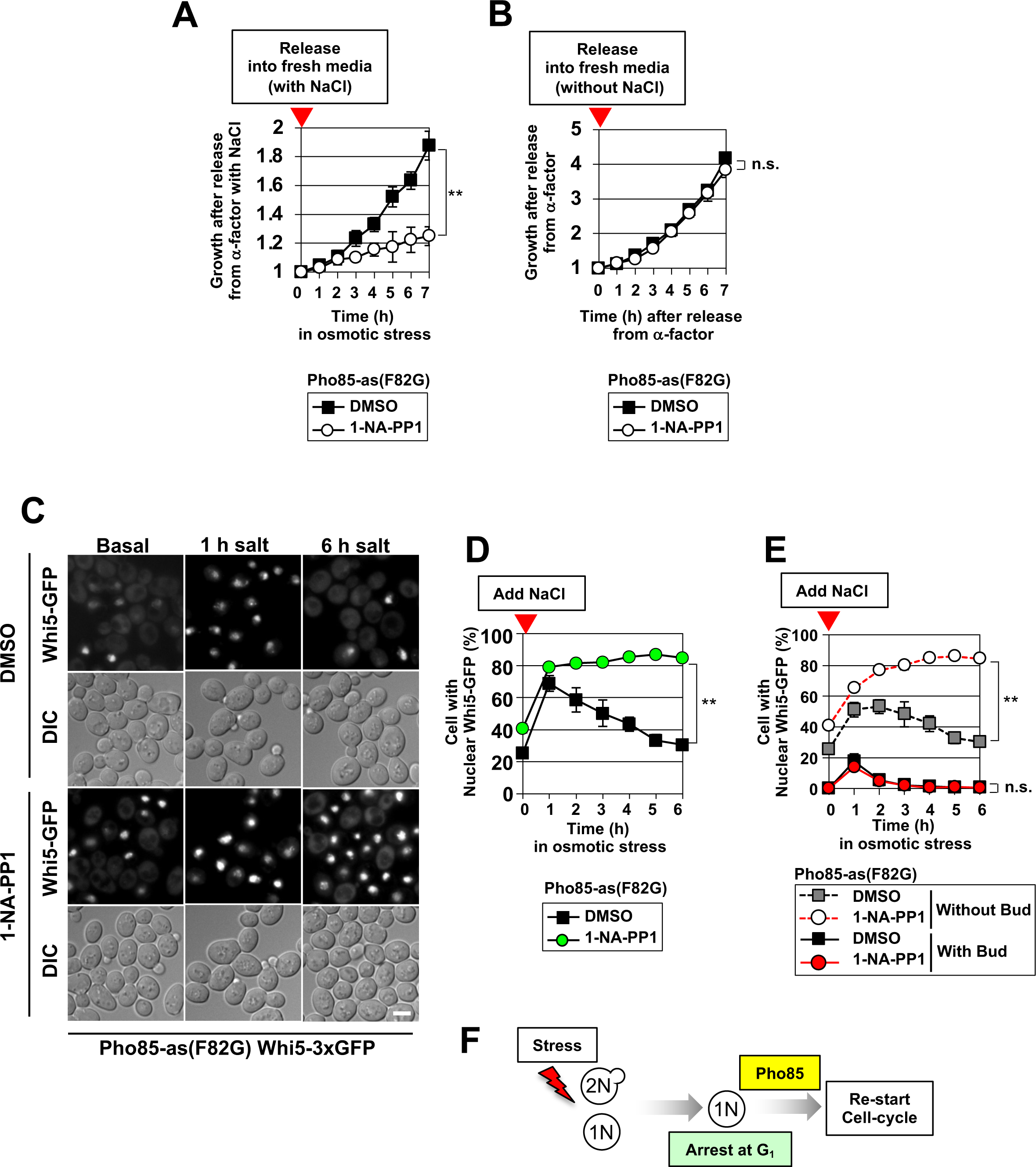
Following hyperosmotic stress, CDK/Pho85 is required to restart the cell-cycle. (A) Pho85 activity is required for growth following hyperosmotic stress, (B) but is not essential for growth under basal conditions. A Pho85-as(F82G) mutant was treated with α-factor plus either DMSO or 20 μM 1-NA-PP1 for 3 h. 80% of the population arrested at G_1_ phase as assessed by DNA content. Cells were released to fresh medium with (A) or without (B) 0.9 M NaCl in the presence of either DMSO or 20 μM 1-NA-PP1 and analyzed every hour for 7 h. Cell number was approximated by measuring OD600. To assess growth, each value was standardized to the value at 0 time. Mean values ± SD. **(p-value < 1 × 10^−3^), n.s. = not significant. (C)-(E) As assessed in an asynchronous population, following hyperosmotic stress, the movement of Whi5-3xGFP to the nucleus is transient. However, during inhibition of Pho85, Whi5-3xGFP remained trapped in the nucleus and cells accumulated in G_1_ phase as determined by the absence of yeast with buds. The Pho85-as (F82G) mutant, expressing Whi5-3xGFP was treated with either DMSO or 20 μM 1-NA-PP1 for 3 h, then exposed to 0.9 M NaCl for 1-6 h. (C) Representative fields of cells. Bar: 5 μm. (D) Quantification of percent cells with Whi5-GFP signal in the nucleus. Mean values ± SD; **(p-value < 1 × 10^−3^); n=3 >300 cells counted per condition, per experiment. (E) Data presented in (D) reporting cells with a bud separately from unbudded cells. After 6 h of treatment with high salt and inhibition of Pho85 activity, Whi5 is trapped in the nucleus of the cells, and most of these cells lack a bud. (F) Model: In hyperosmotic stress, initiation of cell-cycle arrest occurs at any stage, but cells eventually progress to early G_1_ phase prior to the restart of the cell-cycle. CDK/Pho85 is required to restart the cell-cycle from G_1_ phase, following hyperosmotic stress.

Pho85 is an upstream regulator of multiple pathways, thus the *pho85Δ* mutant is sick and exhibits pleiotropic phenotypes. Therefore, we utilized an analog-sensitive point mutant of Pho85, Pho85^F82G^ (Pho85-as) to acutely inhibit Pho85 activity (Carroll et al., 2001). To assess cell-cycle progression following hyperosmotic stress, we monitored the nuclear export/import of Whi5. We found that hyperosmotic stress induced the nuclear import of Whi5 in both budded and unbudded yeast (Supplemental Fig 1D and 1E). This observation fits with an earlier report that hyperosmotic stress causes a transient arrest at multiple phases of the cell-cycle (Argüello-Miranda et al., 2022), and that Whi5 can re-enter the nucleus during stress (Irvali et al., 2023). Note that following stress the nuclear localization of Whi5 was transient. After 6 h of treatment with high salt, Whi5 was exported from the nucleus (Fig 2C). We tested and found that following stress, Pho85 activity is not required for stress-induced import of Whi5 to the nucleus (Fig 2C and 2D). In contrast, after 6 h of treatment with high salt combined with inhibition of Pho85 activity, Whi5 remained in the nucleus (Fig 2C and 2D). These findings show that following stress Pho85 activity is required for the export of Whi5 from the nucleus. Importantly, after 6 h treatment combined with inhibition of Pho85 activity, all cells with Whi5 in the nucleus were unbudded (Fig 2E). This indicates that following stress, in the absence of Pho85 activity, cells remain in G_1_ phase.

Whi5 translocation to the nucleus in stress-induced cell-cycle arrest at G_1_ phase, is likely controlled in part via the Hog1 signaling pathway (González-Novo et al., 2015). However, salt-shock-dependent import of Hog1 into the nucleus is independent of Pho85 activity (Supplemental Fig 1F). Taken together, these findings suggest that (1) in hyperosmotic stress, initiation of cell-cycle arrest occurs at any stage, but cells eventually progress to early G_1_ phase prior to the restart of the cell-cycle; and (2) Pho85 activity is required to restart the cell-cycle from G_1_ phase (Fig 2F), and that Hog1 import into the nucleus does not require Pho85 activity.

To further test the role of Pho85 in the stress-dependent restart of the cell-cycle, we tested the ability of synchronized cells to progress through G_1_ phase during acute inhibition of Pho85. We arrested a Pho85-as mutant expressing Whi5-GFP in early G_1_ phase and then released the cells into high salt. After 1 h of treatment with high salt, Whi5 localized to the nucleus and the cells had 1N DNA content (Fig 3A-C). After longer exposure to hyperosmotic stress, Whi5 was released from the nucleus and <20% of cells had 1N DNA content, indicating that cells progressed to G_2_ -M (Fig 3A-C). However, when Pho85 activity was inhibited, even at 7 h in hyperosmotic stress, Whi5 remained in the nucleus and >80% of cells had 1N DNA content (Fig 3A-C), indicating that cells were trapped at early G_1_ phase. Consistent with these findings, following hyperosmotic stress, inhibition of Pho85 activity caused a defect in cell growth (Fig 2A). In contrast, acute inhibition of Pho85 activity did not affect growth under basal conditions (Fig 2B). Taken together, these findings indicate that Pho85 has critical roles to restart the cell-cycle from G_1_ phase, that are specific to recovery from a stress-induced cell-cycle arrest.

**Figure 3.**
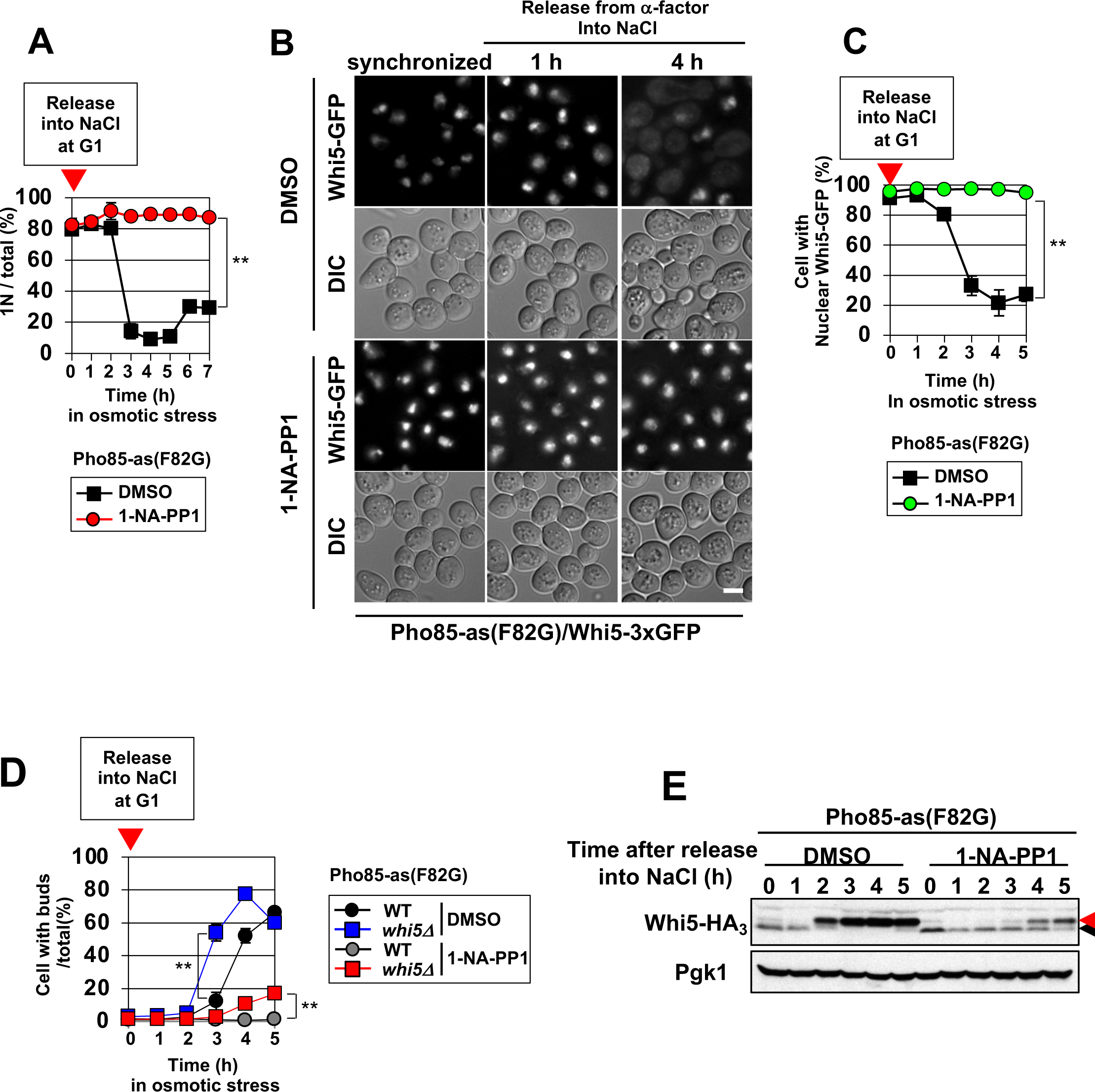
CDK/Pho85 is required to restart the cell-cycle from G_1_ phase following hyperosmotic stress. (A)-(E) The indicated strains were arrested in G_1_ phase with α-factor in the presence of DMSO or 20 μM 1-NA-PP1 for 3 h. Cells were then released to fresh medium with the addition of 0.9 M NaCl, and the continued inclusion of DMSO or 1-NA-PP1, respectively. (A) Following stress, Pho85 activity is required to restart the cell-cycle from G_1_ phase. Following introduction into fresh media, Pho85-(F82G)as cells were analyzed every hour for 7 h. DNA content was measured by flow cytometry. At 0 h, >80% of the cells had 1N DNA content. However, at 3 h following hyperosmotic shock, in the presence of DMSO, only 14% of cells had 1N DNA content. In contrast, the inhibition of Pho85-as by 1-NA-PP1 caused a defect in cell-cycle progression through G_1_ phase where >80% of cells still had 1N DNA. Mean values ± SD; **(p-value < 1 × 10^−3^). (B) and (C) Following stress, Pho85 activity is required for the export of Whi5 at G_1_ phase. Following introduction into fresh media, Pho85-as cells expressing Whi5-3xGFP were analyzed every hour for 5 h. At 0 h, Whi5-3xGFP is localized to the nucleus in more than 90% of the cells. However, at 3 h following hyperosmotic shock, in the presence of DMSO, only 33% of cells have Whi5-3xGFP in the nucleus. In contrast, acute inhibition of Pho85-as by 1-NA-PP1 caused a defect in the export of Whi5-3xGFP from the nucleus. (B) Representative fields of cells. Bar: 5 μm. (C) Quantification of percent cells with Whi5-GFP signal in the nucleus. Mean values ± SD; n=3 >300 cells counted per condition, per experiment. Quantification of percent cells with Whi5-GFP signal in the nucleus. Mean values ± SD; **(p-value < 1 × 10^−3^); n=3 >300 cells counted per condition, per experiment. (D) Following hyperosmotic stress, Pho85 acts in part via Whi5 to restart the cell-cycle. Following release into high salt medium in the presence of DMSO or 1-NA-PP1, the indicated strains were analyzed every hour for 5 h. At 3 h following hyperosmotic shock after release from α-factor, in the presence of DMSO, deletion of *WHI5* caused an acceleration in the bud emergence. Moreover, when deletion of *WHI5* was combined with acute inhibition of Pho85-as using 1-NA-PP1, the defect in bud emergence was partially rescued. Quantification of percent cells with a small bud. Mean values ± SD; **(p-value < 1 × 10^−3^); n=3 >300 cells counted per condition, per experiment. (E) Following hyperosmotic stress, concomitant with the restart of the cell-cycle from G_1_ phase, Whi5 is phosphorylated. However, during inhibition of Pho85, the phosphorylation of Whi5 is defective. The Pho85-as mutant expressing Whi5-HA_3_ was arrested in G_1_ phase with α-factor in the presence of DMSO or 20 μM 1-NA-PP1 for 3 h. Cells were then released to fresh medium with the addition of 0.9 M NaCl, and the continued inclusion of DMSO or 1-NA-PP1, respectively. Following release into fresh medium with high salt, the indicated cells were analyzed every hour for 5 h. Red triangle, the phosphorylated Whi5; black triangle, the non-phosphorylated Whi5. Western blot: anti-HA or anti-Pgk1(control). Blots representative of 3 independent experiments.

### Whi5 determines the timing of the G_1_-S transition in the Pho85-dependent restart of the cell-cycle following stress

Under basal conditions, the G_1_-S transition is accelerated in a *whi5Δ* mutant, which suggests that Whi5 has a role in determining the timing of START and the G_1_-S transition (Costanzo et al., 2004; de Bruin et al., 2004). Therefore, we tested whether Whi5 plays a role in the timing of the restart of the Pho85-dependent cell-cycle under stress conditions. *WHI5* or *whi5Δ* yeast strains were arrested in early G_1_ phase, then released into fresh medium containing high salt. Similar to basal conditions (Costanzo et al., 2004; de Bruin et al., 2004), in high salt when Pho85 was active, deletion of *WHI5* caused an acceleration of bud emergence (Fig 3D). However, following hyperosmotic stress, when Pho85 activity was inhibited, the *WHI5* strain did not resume cell-cycle progression from G_1_ phase as assessed by bud emergence. Moreover, deletion of *WHI5* partially rescued the defect in the bud emergence (Fig 3D). Together, these findings suggest that following hyperosmotic stress Pho85 mediates the restart of the cell-cycle in part by regulation of Whi5, which plays a role in the timing of START and the G_1_-S transition.

We hypothesized that following stress the phosphorylation of Whi5 is regulated by Pho85 during the re-start of the cell-cycle. A Pho85-as yeast strain expressing Whi5-HA_3_ was arrested at early G_1_ phase, then released into fresh medium containing high salt. After 2 h of treatment with high salt in the presence of Pho85 activity, Whi5 was highly phosphorylated as assessed with a gel-shift assay (Fig 3E and Supplemental Fig 2A). Importantly, concomitant with the phosphorylation of Whi5, yeast exited from the stress-induced G_1_ arrest as assessed by DNA content and the export of Whi5 from the nucleus (Fig 3A-C). In contrast, during treatment with high salt combined with inhibition of Pho85 activity, the phosphorylation of Whi5 was impaired (Fig 3E). Taken together, these data suggest that following hyperosmotic stress, Pho85 activity is required to restart the cell-cycle in part via a direct or indirect role of Pho85 in the phosphorylation of Whi5.

### Following hyperosmotic stress, Pho85-Pho80 initiates a cascade to promote the restart of the cell-cycle via direct phosphorylation of an AGC kinase, Sch9

Under basal conditions, Pho85 acts with the cyclin Pcl9 to directly phosphorylate and regulate Whi5, but does not contribute to Whi5 export from the nucleus (Huang et al., 2009). Moreover, Pho85-Pho80 had been identified as a critical mediator of response to stress, although there was no prior evidence that it could function in cell-cycle progression (Huang et al., 2002). For the regulation of Pho85 activity in the restart of the cell-cycle, we reasoned that Pho85 likely functions with the stress-specific cyclin Pho80. Importantly, Pho80 is the only Pho85 cyclin that is essential for growth of yeast on high-salt plates (Jin et al., 2017). We tested and found that Pho85-Pho80 did not phosphorylate Whi5 *in vitro* (Fig 4A). Another way that Pho85 is potentially involved is via action on the Cdc28 inhibitor, Sic1, a key inhibitor of Cdc28, which is phosphorylated by Pho85 upon DNA damage for release from G_1_ arrest (Wysocki et al., 2006) However, we did not pursue this avenue further because in our strain background, the *sic1Δ* mutant had problems emerging from cell-cycle arrest.

**Figure 4.**
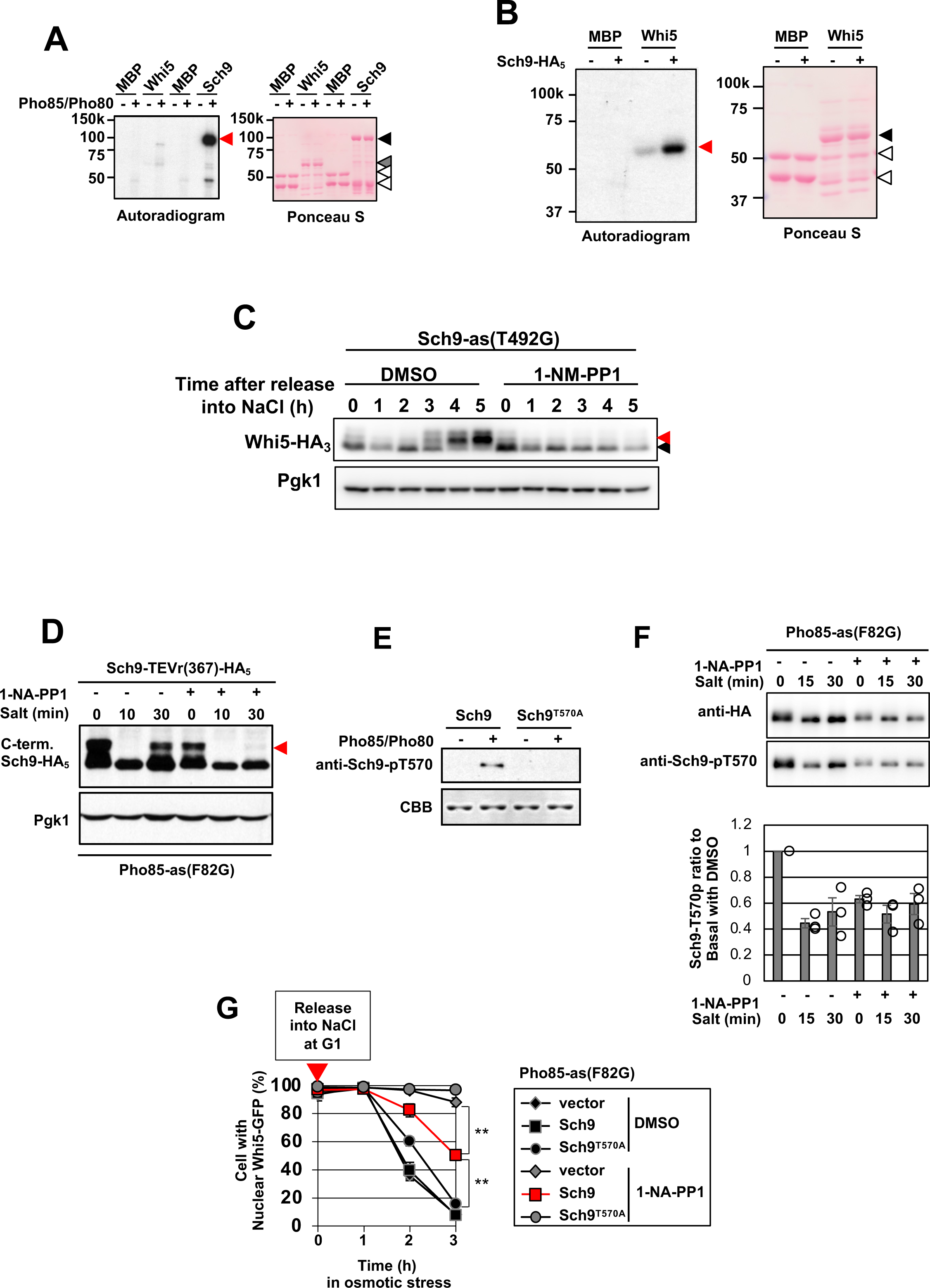
Following hyperosmotic stress, CDK/Pho85 initiates a cascade to restart the cell-cycle via direct phosphorylation and positive regulation of an AGC kinase, Sch9. (A) The CDK Pho85-cyclin Pho80 directly phosphorylates the C terminus of Sch9, but does not phosphorylate Whi5 *in vitro*. pHA-Pho80 was expressed in a *pho80Δ* mutant, immunoprecipitated using anti-HA antibody, and then incubated with [γ-^32^P] ATP and recombinant MBP, MBP-Sch9 (367-824) (see Supplemental Figure 2B), or MBP-Whi5 (full length). Red triangle, phosphorylated C terminus of Sch9; black triangle, MBP-Sch9; gray triangle, MBP-Whi5; white triangles, MBP. Autoradiogram representative of 3 independent experiments. (B) Sch9 directly phosphorylates the C terminus of Whi5 *in vitro*. pSch9-HA_5_ was expressed in a *sch9Δ* mutant, immunoprecipitated using anti-HA antibody, and then incubated with [γ-^32^P] ATP and recombinant MBP or MBP-Whi5 (full length). Red triangle - phosphorylated MBP-Whi5; black triangles - MBP-Whi5; white triangles - MBP. Autoradiogram representative of 3 independent experiments. (C) Following hyperosmotic stress, concomitant with the restart of the cell-cycle from G_1_ phase, Whi5 is phosphorylated. However, during inhibition of Sch9, the phosphorylation of Whi5 is defective. The Sch9-as(T492G) mutant expressing Whi5-HA_3_ was arrested in G_1_ phase with α-factor in the presence of DMSO or 10 μM 1-NM-PP1 for 3 h. Cells were then released to fresh medium with the addition of 0.9 M NaCl, and the continued inclusion of DMSO or 1-NM-PP1, respectively. Following release into fresh medium with high salt, the indicated cells were analyzed every hour for 5 h. Red triangle - phosphorylated Whi5; black triangle - non-phosphorylated Whi5. Western blot: anti-HA or anti-Pgk1(control). Blots representative of 3 independent experiments. (D) Following hyperosmotic stress, Pho85 activity is required for the re-phosphorylation of Sch9. Pho85-as(F82G) mutant expressing pSch9-TEVr(367)-HA_5_ (see supplemental Figure 1D) was treated with DMSO or 20 μM 1-NA-PP1 for 3 h and then treated with 0.9 M NaCl for the times indicated. Cells were lysed with 10% TCA, and digested with TEV protease. Western blot: anti-HA or anti-Pgk1 (control). Blots representative of 3 independent experiments. Red triangle - the phosphorylated C terminus of Sch9. (E) The T570 residue in the consensus kinase activation loop is phosphorylated by the CDK Pho85-cyclin Pho80 *in vitro*. pHA-Pho80 was expressed in a *pho80Δ* mutant, immunoprecipitated using anti-HA antibody and incubated with ATP and MBP-Sch9 (367-824) or MBP-Sch9^T570A^ (367-824). Western blot: anti-phospho Sch9 T570; CBB, loading control. Blot representative of 3 independent experiments. (F) Pho85 activity is required for the phosphorylation of Sch9 T570 under basal condition. Pho85-as(F82G) mutant expressing pSch9-TEVr(367)-HA_5_ was treated with DMSO or 20 μM 1-NA-PP1 for 3 h and then treated with 0.9 M NaCl for the times indicated. Cells were lysed with 10% TCA, and the immunoprecipitation assay of Sch9 was performed using anti-HA. Bound samples were analyzed by SDS-PAGE gel. Western blot: anti-HA or anti-Sch9-pT570. Blots representative of 3 independent experiments. The Sch9 T570 phosphorylation levels were quantified based on the ratio of the signals obtained with anti-p-Sch9^T570^ and anti-HA antibodies and normalized to the ratio obtained for cells treated DMSO under basal condition (time=0 min). (G) Sch9 acts downstream of Pho85 to restart the cell-cycle following hyperosmotic stress. A Pho85-as mutant expressing Whi5-3xGFP with either empty vector, *ADHp*-Sch9 or *ADHp*-Sch9^T570A^ was arrested in G_1_ phase with α-factor in the presence of DMSO or 20 μM 1-NA-PP1 for 3 h. Cells were then released to fresh medium with the addition of 0.9 M NaCl, and the continued inclusion of DMSO or 1-NA-PP1, respectively. Following release into fresh media, cells were analyzed every hour for 3 h. At 0 h, Whi5-3xGFP was localized to the nucleus in more than 90% of the cells. However, by 3 h following hyperosmotic shock, in the presence of DMSO, with either empty vector, over expression of Sch9 or Sch9^T570A^, Whi5-3xGFP was exported from the nucleus. However, during inhibition of Pho85 by 1-NA-PP1, there was a defect in the export of Whi5-3xGFP from the nucleus which was not rescued by overexpression of empty vector or Sch9^T570A^. However, overexpression of wild-type Sch9 rescued the defect in the export of Whi5 from the nucleus. Quantification of percent cells with Whi5-GFP signal in the nucleus. Mean values ± SD; **(p-value < 1 × 10^−3^); n=3 >300 cells counted per condition, per experiment.

To gain mechanistic insights into how Pho85-Pho80 restarts the cell-cycle following stress, we sought to determine the direct target of Pho85 in the regulation of Whi5. In considering possible targets of Pho85, we tested an AGC kinase, Sch9. Sch9 signaling from the vacuole/lysosome is essential for cell-cycle progression from early G_1_ phase (Jin et al., 2022; Jin and Weisman, 2015). Moreover, under basal conditions Pho85-Pho80 is required for the phosphorylation of Sch9 to prime Sch9 for subsequent phosphorylation and activation by TORC1 (Deprez et al., 2023). Importantly, the C terminus of Sch9 is acutely dephosphorylated and then re-phosphorylated during many types of stress, including hyperosmotic stress (Urban et al., 2007). However, the kinases that re-phosphorylate Sch9 following stress were unknown. Thus, we tested whether Sch9 is directly phosphorylated by Pho85-Pho80. We found that an immunoprecipitate of Pho85-Pho80 from yeast lysates directly phosphorylates a recombinant Sch9 peptide (367-824 amino acids) *in vitro* (Fig 4A and Supplemental Fig 2B). In addition, we found that Sch9 directly phosphorylates Whi5 *in vitro* (Fig 4B), which indicates that Whi5 is a direct downstream target of Sch9. Note that in this experiment, only phosphorylated Whi5 is detected *in vitro*. To test whether Sch9 is critical for phosphorylation of Whi5 *in vivo*, we utilized Sch9^T492G^(Sch9-as), an analogue sensitive mutant (Plank et al., 2020), to acutely inhibit Sch9. Importantly, following hyperosmotic stress, Sch9 activity is required for the phosphorylation of Whi5 during cell-cycle progression (Fig 4C). These observations suggest that Sch9 is critical for the phosphorylation of Whi5 following hyperosmotic stress.

We next tested whether Pho85 activity is required *in vivo* for the activation of Sch9 during hyperosmotic stress. As previously reported (Urban et al., 2007a), the C terminus of Sch9 is highly phosphorylated in a TORC1-dependent manner (Supplemental Fig 2C) and the phosphorylation of Sch9 is required for Sch9 activation. To monitor the phosphorylation status of the C terminus of Sch9, we engineered a TEV cleavage site (TEVr) after Sch9 I366: Sch9-TEVr(367) and included HA_5_ at the C terminus (Supplemental Fig 2B). Consistent with previous observations, when yeast expressing Sch9-TEVr(367)-HA were exposed to hyperosmotic stress, the C terminus of Sch9 was acutely dephosphorylated at 10 min and then re-phosphorylated after 30 min (Fig 4D) (Urban et al., 2007). In contrast, inhibition of Pho85 activity resulted in a defect in the re-phosphorylation of the C terminus of Sch9 following stress (Fig 4D). These observations suggest that Pho85 is critical for the re-phosphorylation of Sch9 following hyperosmotic stress.

Activation of the AGC kinase family including Sch9 requires phosphorylation of a Ser/Thr residue in its activation loop (Jacinto and Lorberg, 2008). Under basal conditions, Pkh1 and Pkh2 directly phosphorylate T570 in the consensus activation loop of Sch9 which is located on its C-terminus (Urban et al., 2007). Notably, Pho85-Pho80 directly phosphorylates Sch9 T570 *in vitro* (Fig 4E). While this is not a canonical CDK T/P site, CDKs use both canonical and non-canonical sites (Jin et al., 2022; Liu et al., 2009; Suzuki et al., 2015). Moreover, we found that under basal conditions, Pho85 activity is required for the phosphorylation of Sch9 T570 *in vivo* (Fig 4F). However, even in the absence of Pho85 inhibition, there was only a partial recovery of phosphorylation at T750 at 30 min following hyperosmotic stress. In contrast, re-phosphorylation of Sch9 as assessed by gel shift fully returns, and these sites either directly, or indirectly require Pho85 activity as well. Importantly, using a more sensitive assay, we found that following stress the phosphorylation of Sch9 T570 is critical to restart the cell-cycle. Pho85-as yeast strains that overexpressed either wild-type Sch9, Sch9^T570A^ or empty vector, were synchronized at early G_1_ phase, then released into fresh medium with high salt. At 3 h following hyperosmotic stress, in the presence of Pho85 activity and overexpression of either empty vector, Sch9 or Sch9^T570A^, Whi5 was released from the nucleus (Fig 4G). Importantly, during inhibition of Pho85 activity, overexpression of wild-type Sch9 partially rescued the defect in the export of Whi5 from the nucleus, whereas overexpression of Sch9^T570A^ or empty vector did not rescue (Fig 4G). These findings indicate that Pho85-dependent restart of the cell-cycle acts in part via its direct phosphorylation of Sch9 at sites which include T570, and likely also includes other sites. It is also possible, that in addition, Pho85 activates additional kinases that also re-phosphorylate Sch9. Together, these findings reveal a pathway the restarts the cell-cycle following stress and show that Pho85-Pho80 directly phosphorylates and activates Sch9, which then directly phosphorylates Whi5, and releases Whi5-dependent inhibition of the cell-cycle at G_1_ phase.

### Pho85 is required to restart the cell-cycle during recovery from nutrient starvation

There are many conditions where cells need to restart the cell-cycle. The finding that Pho85 is critical for the restart of the cell-cycle following hyperosmotic stress, raises the question of whether this Pho85-dependent pathway restarts the cell-cycle following other types of stress. Rapamycin treatment, a mimic of nitrogen starvation, results in an arrest in the cell-cycle, and a concomitant loss of the G_1_-specific Cdc28 cyclins, Cln1 and Cln3 (Barbet et al., 1996). Thus, we tested whether Pho85 is required to restart the cell-cycle during recovery after nitrogen starvation. A Pho85-as mutant expressing Whi5-GFP was incubated with nitrogen starvation medium and then re-fed normal nutrient-rich medium. Under starvation conditions, Whi5 localized to the nucleus in the presence or absence of Pho85 activity (Fig 5A and 5B). Following recovery for 1 h in nutrient-rich medium, when Pho85 was active, Whi5 was exported from the nucleus (Fig 5B and 5C). In contrast, inhibition of Pho85 activity delayed the export of Whi5 from the nucleus (Fig 5B and 5C). This suggests that Pho85 is likely required for cell-cycle progression during recovery from nitrogen starvation. As a further test, we arrested the Pho85-as mutant at early G_1_ phase and then released the yeast into starvation medium for 3 h in the presence or absence of Pho85 inhibition. Then yeast cells were re-fed with nutrient medium. Whi5 localization as well as DNA content and growth rate were assessed (Fig 5D-G and Supplemental Fig 2D). We found that inhibition of Pho85 activity caused a growth defect during recovery from nutrient starvation (Fig 5E). Moreover, inhibition of Pho85 activity also caused a block in the export of Whi5 from the nucleus, as well as a block in cell-cycle progression through G_1_ phase as assessed by DNA content (Fig 5F, 5G and Supplemental Fig 2D). During recovery from nutrient starvation, the expression of Cdc28-cyclins was transiently downregulated (Supplemental Fig 2E). Notably, the re-expression of Cln1, Cln2 and Clb6 occurred concomitant with the rephosphorylation of Whi5 (Fig 5H and Supplemental Fig 2E). Note that, when Pho85 activity was inhibited during recovery from nutrient starvation, Whi5 phosphorylation did not occur (Fig 5H).

**Figure 5.**
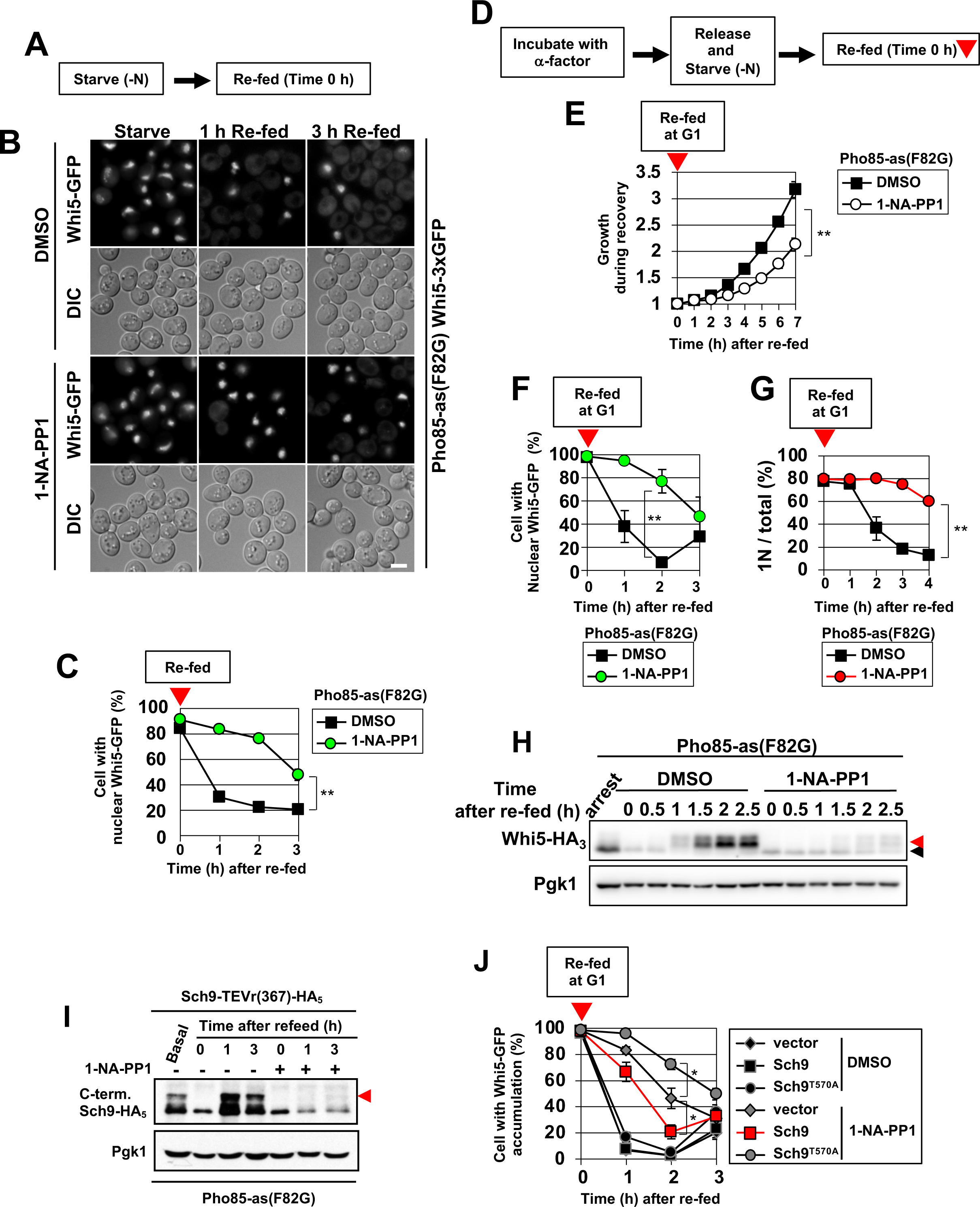
CDK Pho85 is required to restart the cell-cycle during recovery following nitrogen starvation. (A) Schematic of experiment for (B) and (C). Acute inhibition of Pho85 causes a defect in the export of Whi5-3xGFP from the nucleus following recovery after nitrogen starvation. A Pho85-as(F82G) mutant expressing Whi5-3xGFP cultured in nitrogen starvation medium with DMSO or 20 μM 1-NA-PP1 for 3 h and then released to nutrient-rich medium in the presence of DMSO or 20 μM 1-NA-PP1 for 1-3 h. (B) Representative fields of cells. Bar: 5 μm. (C) Quantification of percent cells with accumulated Whi5-GFP signal in the nucleus. Mean values ± SD; **(p-value < 1 × 10^−3^); n=3 >300 cells counted per condition, per experiment. (D) Schematic of experiment for (E)-(H). The indicated strains were arrested in G_1_ phase with α-factor in the presence of DMSO or 20 μM 1-NA-PP1 for 3 h. Cells were released into nitrogen starvation medium for 3 h and then introduced into nutrient-rich medium. DMSO or 20 μM 1-NA-PP1, as indicated, were present during the arrest, release into starvation medium and introduction into nutrient-rich medium. (E) Pho85 activity is required for growth from G_1_ phase, following nitrogen starvation. Following re-feeding with nutrient-rich medium with DMSO or 20 μM 1-NA-PP1 for 1-7 h, cell number was approximated using OD600. To report growth, each value was divided by the value at 0 time. Mean values ± SD; **(p-value < 1 × 10^−3^). (F) Acute inhibition of Pho85 causes a defect in the export of Whi5 from the nucleus at G_1_ phase following recovery after nitrogen starvation. Following re-feeding with nutrient-rich medium with DMSO or 20 μM 1-NA-PP1 for 1-3 h, Whi5 localization was determined in a Pho85-as(F82G) mutant expressing Whi5-3xGFP. Quantification of percent cells with accumulated Whi5-GFP signal in the nucleus. Mean values ± SD; **(p-value < 1 × 10^−3^); n=3 >300 cells counted per condition, per experiment. (G) Pho85 activity is required to restart the cell-cycle from G_1_ phase following recovery after nitrogen starvation. Following re-feeding with nutrient-rich medium with DMSO or 20 μM 1-NA-PP1 for 1-4 h, DNA content was measured in the Pho85-as(F82G) mutant using flow cytometry. Mean values ± SD; **(p-value < 1 × 10^−3^); n=3. (H) Following recovery after nitrogen starvation, concomitant with its nuclear export, Whi5 is phosphorylated. However, during inhibition of Pho85, the phosphorylation of Whi5 is defective. Red triangle -phosphorylated Whi5; black triangle - non-phosphorylated Whi5. Western blot: anti-HA or anti-Pgk1(control). Blots representative of 3 independent experiments. (I) During recovery after nitrogen starvation, Pho85 activity is required for the re-phosphorylation of Sch9. Pho85-as(F82G) containing cells expressing pSch9-TEVr(367)-HA_5_ were cultured in the nitrogen starvation medium for 3 h in the presence of either DMSO or 20 μM 1-NA-PP1. Then cells were released into nutrient-rich medium. DMSO or 20 μM 1-NA-PP1, as indicated, were present during culture in the starvation medium and introduction into nutrient-rich medium. Cells were lysed with 10% TCA, and digested with TEV protease. Western blot: anti-HA or anti-Pgk1 (control). Blots representative of 3 independent experiments. Red triangle, phosphorylated C terminus of Sch9. (J) Sch9 acts downstream of Pho85 to restart the cell-cycle during recovery after nitrogen starvation. An empty vector, Sch9 or Sch9^T570A^ were overexpressed in a Pho85-as(F82G) mutant. At the 2 h time point during recovery after nitrogen starvation, in the presence of DMSO, with either empty vector, or overexpression of Sch9 or Sch9^T570A^, Whi5-3xGFP was exported from the nucleus. In contrast, during inhibition of Pho85 by 1-NA-PP1, either empty vector or overexpression of Sch9^T570A^ causes a defect in the export of Whi5-3xGFP from the nucleus. However, overexpression of Sch9 rescued the defect in the export of Whi5 from the nucleus. Quantification of percent cells with Whi5-GFP signal in the nucleus. Mean values ± SD; *(p-value < 3 × 10^−3^); n=3 >300 cells counted per condition, per experiment.

To assess whether Pho85-Pho80 restarts the cell-cycle during recovery from nutrient starvation via the same pathway used to restart of the cell-cycle during hyperosmotic stress, we tested whether Pho85 is required for nutrient-dependent re-phosphorylation of Sch9. The C terminus of Sch9 is acutely dephosphorylated and then re-phosphorylated during recovery from nutrient starvation (Urban et al., 2007). Consistent with this previous finding, when Pho85-as yeast expressing Sch9-TEVr(367)-HA_3_ were starved, the C terminus of Sch9 was acutely dephosphorylated and then re-phosphorylated within 1 h of refeeding (Fig 5I). However, when Pho85 was inhibited during recovery from starvation, re-phosphorylation of the C terminus of Sch9 did not occur (Fig 5I). Furthermore, we found that during recovery from starvation, phosphorylation of Sch9 on T570 was critical to restart the cell-cycle. Pho85-as yeast strains expressing Whi5-GFP that overexpressed either wild-type Sch9, Sch9^T570A^ or empty vector were arrested in G_1_ phase, released into starvation medium for 3 h, then re-fed with nutrient medium. In the presence of Pho85 activity with overexpression of either empty vector, Sch9 or Sch9^T570A^, after 3 h of re-feeding, Whi5 was released from the nucleus (Fig 5J). In contrast, when Pho85 was inhibited, overexpression of wild-type Sch9, but not empty vector or the Sch9^T570A^ mutant, rescued the defect in the export of Whi5 from the nucleus (Fig 5J).

### During recovery from stress, Pho85 and Cdc28 function in parallel to restart the cell-cycle

We tested whether following stress the new synthesis of each cyclin requires Pho85 activity. Notably, we found that Pho85 activity is required for the expression of some but not all of the cyclins. Cells were arrested in early G_1_ phase, then released into fresh medium containing high salt. In high salt when Pho85 was active, the expression of Cln1; a G_1_-specific cyclin, Clb6; a G_1_-S specific cyclin or Clb2; a G_2_-M specific cyclin resumed, which would allow the restart and progression of cell-cycle. However, when Pho85 activity was inhibited, Cln1 was re-expressed and the expression of Clb6 and Clb2 was absent (Fig 6A). This data raised the possibility that Cdc28 is also required for G_1_ progression during hyperosmotic stress. We utilized the Cdc28-as1 mutant and found that when Cdc28 activity was inhibited the phosphorylation of Whi5 did not occur (Fig 6B). Whi5 contains 12 putative CDK1/Cdc28 phosphorylation sites plus 6 non-CDK1/Cdc28 sites which are phosphorylated *in vivo* (Wagner et al., 2009). We found that the 12 known CDK1/Cdc28 phosphorylation sites on Whi5, but not the non-CDK1/Cdc28 sites are required for the translocation of Whi5 from the nucleus during recovery from hyperosmotic stress (Supplemental Fig 2F). Taken together, our findings suggest that (1) the non-canonical CDK Pho85 is critical to restart the cell-cycle in at least two types of stress and (2) following stress Pho85-Sch9 signaling positively regulates the progression of the cell-cycle through G_1_ phase via direct phosphorylation of Sch9, which in turn directly phosphorylates Whi5, and (3) Pho85 acts in parallel with Cdc28 to re-start the cell-cycle via a parallel regulation of progression through G_1_-phase (Fig 6C).

**Figure 6.**
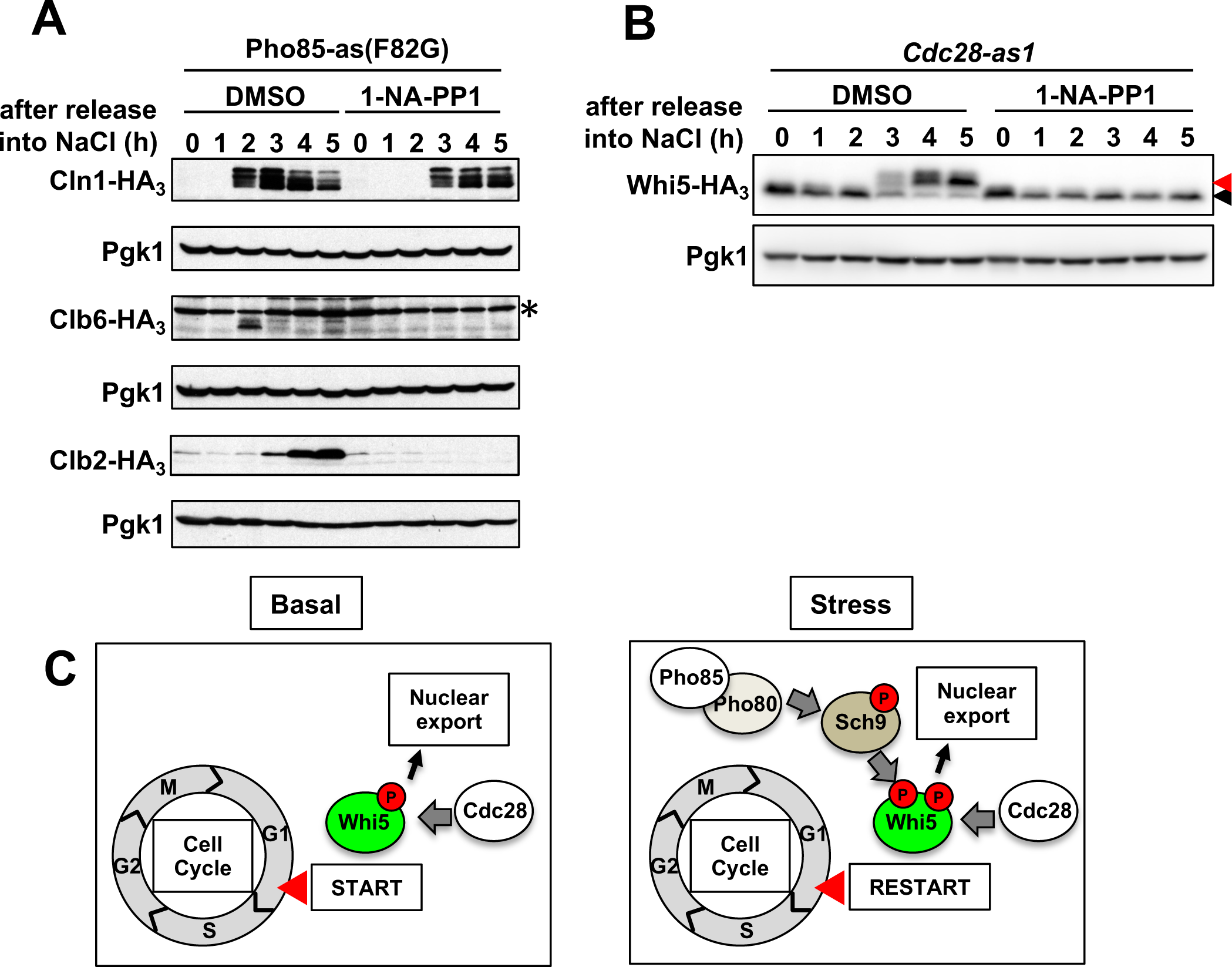
During stress, Pho85 functions in parallel with Cdc28 to restart the cell-cycle and progression through G_1_-phase. (A) Following hyperosmotic stress, concomitant with the restart of the cell-cycle from G_1_ phase, Pho85 activity is required for the G_1_/S transition in part via the expression of the Cdc28/CDK1 S-phase specific cyclin, Clb6. The Pho85-as(F82G) mutant expressing Cln1-HA_3_, Clb6-HA_3_ or Clb2-HA_3_ were arrested in G_1_ phase with α-factor in the presence of DMSO or 20 μM 1-NA-PP1 for 3 h. Cells were then released to fresh medium with the addition of 0.9 M NaCl, and the continued inclusion of DMSO or 1-NA-PP1, respectively. Following release into fresh medium with high salt, the indicated cells were analyzed every hour for 5 h. Western blot: anti-HA or anti-Pgk1(control). Blots representative of 3 independent experiments. Black asterisk indicates background band. (B) Following hyperosmotic stress, concomitant with the restart of the cell-cycle from G_1_ phase, Cdc28 function is required for the phosphorylation of Whi5. The Cdc28-as1 mutant expressing Whi5-HA_3_ was arrested in G_1_ phase with α-factor in the presence of DMSO or 10 μM 1-NM-PP1 for 3 h. Cells were then released to fresh medium with the addition of 0.9 M NaCl, and the continued inclusion of DMSO or 1-NM-PP1, respectively. Following release into fresh medium with high salt, the indicated cells were analyzed every hour for 5 h. Red triangle - phosphorylated Whi5; black triangle - non-phosphorylated Whi5. Western blot: anti-HA or anti-Pgk1(control). Blots representative of 3 independent experiments. (C) Model: Under basal conditions, cell-cycle progression from early G_1_ (START) is achieved via CDK1/Cdc28 which phosphorylates a few key substrates includingWhi5. Whi5 is exported from the nucleus. Following stress, a non-canonical CDK and cyclin, CDK5/Pho85-cyclin Pho80, directly phosphorylates Sch9. Then activated Sch9 directly phosphorylates Whi5, which is critical for the restart of the cell-cycle from G_1_ phase. It is possible that Pho85-Pho80 phosphorylates additional targets as well. In parallel with Pho85-Pho80, CDK1/Cdc28 also functions in the phosphorylation of Whi5 following stress. Moreover, the export of Whi5 from the nucleus contributes to the transition from G_1_ to S phase via the expression of S-phase specific cyclins for CDK1/Cdc28.

## Discussion

Our studies reveal that there are multiple differences between regulation of the cell cycle under basal conditions, vs. regulation of the cell cycle during stress. Of particular note, we observed that Cln3, the earliest CDK1/Cdc28 cyclin for G_1_ progression, is downregulated during stress and surprisingly Cln3 levels are barely detectable after the cell-cycle resumes. These discoveries strongly suggest that additional pathways restart the cell-cycle and may potentially act in parallel with CDK1/Cdc28.

Our studies show that following stress, the re-start of the cell-cycle at G_1_-phase is regulated by the non-canonical CDK, Pho85 and that Pho85 co-operates with the canonical CDK, Cdc28 (Fig 6C). We find that Sch9 is a direct target of Pho85 and that active Sch9 directly phosphorylates Whi5 which promotes the G_1_-S transition. Following many types of stresses, the phosphorylation of Whi5 by Sch9 likely induces the export of Whi5 from the nucleus during the restart the cell-cycle from G_1_ phase. This contrasts with basal conditions, where Cdc28-Cln3 plays an essential role in cell-cycle progression from early G_1_ phase (Costanzo et al., 2004; de Bruin et al., 2004; Litsios et al., 2022; Schmoller et al., 2015). It is tempting to speculate that even during basal conditions, Sch9 may directly phosphorylate Whi5. In basal conditions the direct activation of Sch9 occurs via TORC1 and the non-canonical CDK, Bur1-Bur2 (Jin et al., 2022; Jin and Weisman, 2015), and interestingly, Pho85 also primes Sch9 to be a substrate of TORC1 (Deprez et al., 2023). Moreover, in basal conditions, a START repressor, Whi7 is phosphorylated by Pho85 (Ros-Carrero et al., 2024). Together, Pho85 plays multiple roles in the cell cycle, both under basal conditions and during stress.

We also uncovered additional stress-dependent changes in Whi5 that are independent of Pho85-Pho80. In the canonical cell-cycle, the import of Whi5 to the nucleus is confined to late metaphase. However, we and others found that following stress, Whi5 can be imported into the nucleus at multiple phases of the cell-cycle (Supplemental Fig 1D) and (Irvali et al., 2023).

The type of signal that activates Pho85-Pho80 following stress remains unknown. Moreover, the signal that restarts the cell-cycle following hyperosmotic stress may be distinct from the signal that restarts the cell-cycle following refeeding. However, it is notable that the yeast vacuole plays critical roles both for response to changes in osmolarity, and for sensing nutrient availability. This raises the possibility that the vacuole plays a role in the activation of Pho85-Pho80 under both conditions. In further support of this hypothesis, during stress Pho85 has a second role at the vacuole/lysosome and acts directly on the Fab1 complex (Jin et al., 2017). During hyperosmotic stress, Pho85-Pho80 activation of Fab1 regulates a large transient increase in the signaling lipid, phosphatidylinositol (3,5)-bisphosphate, which provides early protection for yeast viability prior to the establishment of adaptation pathways that signal from the nucleus (Jin et al., 2017). That Pho85-Pho80 phosphorylates Fab1 on the vacuole, indicates that during stress, a pool of Pho85-Pho80 is present on the vacuole.

Since virtually all of the cellular pool of Pho80 remains in the nucleus, the hypothesis that Pho85-Pho80 restarts the cell-cycle by signaling from the vacuole could not be tested further (Jin et al., 2017). It is tempting to speculate that following stress, Pho85 may regulate additional stress-related responses and may serve as a hub on the vacuole for stress-related pathways.

It is notable that yeast utilize an extreme situation, transient shutdown of Cdc28/CDK1, to pause cell-cycle progression. This finding further underscores the hypothesis that G_1_ to S progression and DNA synthesis need to be postponed until stressful conditions are mitigated.

Our discovery of a parallel pathway for the canonical yeast Cdc28/CDK1 cell-cycle raises the possibility that mammalian cells utilize similar strategies. For example, stem cells need to maintain a non-proliferative state and under specific stimuli re-enter the cell-cycle to proliferate. In addition, cancer stem cells maintain a non-proliferative state for many years and may use a similar parallel pathway to re-enter the cell-cycle (Takeishi and Nakayama, 2016; Yoshida and Saya, 2016). One mammalian cell candidate for a parallel cell-cycle pathway is CDK5, the mammalian homolog of Pho85. This possibility fits with our earlier finding that Pho85 and CDK5 each phosphorylate Fab1 and PIKfyve, respectively, and in that situation, act as true homologs. Future studies of cell-cycle pathways that act in parallel with canonical cell-cycle CDK kinases, may provide new therapeutics for human diseases including cancer.

## Material and Methods

### Strains, plasmids and media

Yeast strains and plasmids are listed in Tables 1 and 2. Yeast grown in YEPD (1% yeast extract, 2% peptone, 2% dextrose), synthetic complete (SC) medium (containing 2% dextrose and lacking selective supplement(s)) or starvation medium (lacking ammonium sulfate and containing 2% dextrose), at 24°C. For synchronization of yeast at early G_1_ phase, cells were treated with 3.7 μM α-factor (Zymo Research) or 9 μM α-mating factor (Peptide Institute, Inc.) for three hours, washed twice, then released to the indicated media. For acute inhibition of Pho85 activity, Pho85-as(F82G) mutant was treated with 20 μM 1-NA-PP1 (Cayman) in DMSO for three hours, then assessed. For acute inhibition of Sch9 activity, the Sch9-as(T492G) mutant was treated with 10 μM 1-NM-PP1 (Cayman) in DMSO for three hours, then assessed. For acute inhibition of Cdc28 activity, the Cdc28-as1 mutant (Bishop et al., 2000) was treated with 10 μM 1-NM-PP1 (Cayman) in DMSO for three hours, then assessed.

**Table 1.** Yeast strains used in this study.

| Strain | Genotype | Source | Figure |
| --- | --- | --- | --- |
| LWY7235 | *MATa, ura3-52, leu2-3,-112, his3-Δ200, trp1-Δ901, lys2-801, suc2-Δ9* | (Bonangelino et al., 1997, p. 7) | 1, S1,2 |
| NJ73 | *MATa, ura3-52, leu2-3,-112, his3-Δ200, trp1-Δ901, lys2-801, suc2-Δ9, Cln1-HA<sub>3</sub>::HIS3* | This study | 1, S1,2 |
| NJ75 | *MATa, ura3-52, leu2-3,-112, his3-Δ200, trp1-Δ901, lys2-801, suc2-Δ9, Cln2-HA<sub>3</sub>::HIS3* | This study | 1, S1,2 |
| NJ77 | *MATa, ura3-52, leu2-3,-112, his3-Δ200, trp1-Δ901, lys2-801, suc2-Δ9, Clb1-HA<sub>3</sub>::HIS3* | This study | 1, S1,2 |
| NJ79 | *MATa, ura3-52, leu2-3,-112, his3-Δ200, trp1-Δ901, lys2-801, suc2-Δ9, Clb2-HA<sub>3</sub>::HIS3* | This study | 1, S1,2 |
| NJ85 | *MATa, ura3-52, leu2-3,-112, his3-Δ200, trp1-Δ901, lys2-801, suc2-Δ9, Clb5-HA<sub>3</sub>::HIS3* | This study | 1, S1,2 |
| NJ87 | *MATa, ura3-52, leu2-3,-112, his3-Δ200, trp1-Δ901, lys2-801, suc2-Δ9, Clb6-HA<sub>3</sub>::HIS3* | This study | 1, S1,2 |
| NJY879 | *MATa, ura3-52, leu2-3,-112, his3-Δ200, trp1-Δ901, lys2-801, suc2-Δ9, Cln3-HA<sub>3</sub>::HIS3* | This study | 1, S1,2 |
| LWY17519 | *MATa, ura3-52, leu2-3,-112, his3-Δ200, trp1-Δ901, lys2-801, suc2-Δ9, Whi5-3xGFP::URA3* | This study | S1 |
| LWY17518 | *MATa, ura3-52, leu2-3,-112, his3-Δ200, trp1-Δ901, lys2-801, suc2-Δ9, Whi5-3xGFP::URA3, Pho85F82G::Nat* | This study | 2-5, S2 |
| LWY16217 | *MATa, ura3-52, leu2-3,-112, his3-Δ200, trp1-Δ901, lys2-801, suc2-Δ9, Pho85F82G::Nat* | This study | 2-5, S1 |
| NJ104 | *MATa, ura3-52, leu2-3,-112, his3-Δ200, trp1-Δ901, lys2-801, suc2-Δ9, Cln1-HA<sub>3</sub>::HIS3, Pho85F82G::Nat* | This study | 6 |
| NJ161 | *MATa, ura3-52, leu2-3,-112, his3-Δ200, trp1-Δ901, lys2-801, suc2-Δ9, Clb6-HA<sub>3</sub>::HIS3, Pho85F82G::Nat* | This study | 6 |
| NJ145 | *MATa, ura3-52, leu2-3,-112, his3-Δ200, trp1-Δ901, lys2-801, suc2-Δ9, Clb2-HA<sub>3</sub>::HIS3, Pho85F82G::Nat* | This study | 6 |
| LWY14347 | *MATa, ura3-52, leu2-3,-112, his3-Δ200, trp1-Δ901, lys2-801, suc2-Δ9, sch9Δ::KanMX, pRS416 SCH9* |  | 4 |
| NJ303 | *MATa, ura3-52, leu2-3,-112, his3-Δ200, trp1-Δ901, lys2-801, suc2-Δ9, Whi5-HA<sub>3</sub>::HIS3, Pho85F82G::Nat* | This study | 3,5, S2 |
| NJ293 | *MATa, ura3-52, leu2-3,-112, his3-Δ200, trp1-Δ901, lys2-801, suc2-Δ9, whi5Δ::KanMX, Pho85F82G::Nat* | This study | 3 |
| NJYJ66 | *MATa, ura3-52::Sch9-TEVr-HA5::URA3, leu2-3,-112, his3-Δ200, trp1-Δ901, lys2-801, suc2-Δ9, Pho85F82G::Nat* | This study | 4 |
| NJYJ92 | *MATa, ura3-52, leu2-3,-112, his3-Δ200, trp1-Δ901, lys2-801, suc2-Δ9, sch9Δ::KanMX, pRS416 SCH9T492G, Whi5-HA<sub>3</sub>::HIS3* | This study | 4 |
| NJYJ149 | *ura3-1 trp1-1 leu2-3, 112 his3-11 ade2-1 can1-100 GAL+ cdc28d::cdc28-as1 bar1::HISG, Whi5-HA<sub>3</sub>::HIS3* | This study | 6 |

**Table 2.** Plasmids used in this study.

| Plasmid name | Description | Source | Figure |
| --- | --- | --- | --- |
| pMAL-Sch9(367-824) | Amp | This study | 4 |
| pMAL- Sch9<sup>T570A</sup>(367-824) | Amp | This study | 4 |
| pMAL-Whi5 | Amp | This study | 4 |
| pVT102-H | 2μ, *HIS3* | (Vernet et al., 1987) | 4,5 |
| pVT102-H *SCH9* | 2μ, *HIS3* | (Jin and Weisman, 2015b) | 4,5 |
| pVT102-H *SCH9*<sup>T570A</sup> | 2μ, *HIS3* | This study | 4,5 |
| pRS413 *Sch9-HA<sub>5</sub>* | CEN, *HIS3* | (Urban et al., 2007b) | 4,S2 |
| pRS413 *Sch9-TEVr(367)-HA<sub>5</sub>* | CEN, *HIS3* | This study | 4,5,S2 |

### Fluorescence Microscopy

Images were acquired using the DeltaVision-Elite Restoration Microscopy System (Applied Precision, Issaquah, WA, USA). For the time course study (Supplemental Fig 2D), cells were attached to slides treated with 1 mg/ml concanavalin A.

### Flow cytometry analysis

FACS analysis of DNA content was previously described (Jin et al., 2022). Briefly, cells were fixed with 70% ethanol, then washed twice with wash buffer (50 mM Tris-HCl (pH7.5)) and treated with 2 mg/ml RNaseA (Sigma-Aldrich) in wash buffer at 37°C overnight. Cells were then treated with 5 mg/ml pepsin for 30 min at room temperature and stained with 50 mg/ml propidium iodide (Sigma-Aldrich) in staining buffer (180 mM Tris-HCl (pH7.5), 180 mM NaCl, 70 mM MgCl_2_) for 1 hour at room temperature. Cells were analyzed by FACS analysis (Fortessa, BD). Approximately, 13,000∼10,000 cells were examined. Representative histograms were shown in Supplemental Fig 3.

### Western blot analysis

Cells were lysed with 10 % TCA and resuspended in 2X SDS sample buffer. Protein extracts were subjected to SDS-PAGE followed by immunoblotting with the indicated antibodies.

### Antibodies for western blot analysis

anti-Pgk1; 1:80,000 (Invitrogen), anti-HA; 1:10,000 (Covance), anti-HA(3F10); 1:3,000 (Sigma-Aldrich), anti-Phospho-Sch9(T570); 1:15,000 (Fig 4E) or 1:5,000 (Fig 4F) (a kind gift from Dr. Loewith, University of Geneva). Blots were visualized using X-lay film, FUSION-FX7 (Viber-Lourmat) imaging system or ImageQuant LAS 500 (Cytiva) imaging system, respectively.

### TEV protease cleavage assay

Mid-log phase cells were lysed with 10 % TCA and resuspended in 2X SDS sample buffer. The dilution buffer (50 mM Tris-HCl (pH 7.5), 150 mM NaCl, 10 μM EDTA, 0.5 % Tween 20) was added to the lysate and incubated with 150 U/mL TEVp (Promega) at 30°C for 1 hour. Reactions terminated with 10 % TCA.

### Protein purification

Proteins expressed in *E. coli* BL21 star (DE3) with pRARE. Cells grown in TB with 0.05 mg/ml ampicillin and 0.025 mg/ml spectinomycin at 37°C to an OD_600_ of 0.3. After 18 hours of induction with 0.2 mM IPTG at 16°C, cells were harvested, sonicated in lysis buffer (20 mM Tris-HCl (pH 7.5), 200 mM NaCl, 1 mM DTT, 1x EDTA-free protease inhibitor (Roche)). Proteins purified with amylose resin (NEB) and dialyzed overnight in Pho85-Pho80 dialysis buffer (10 mM Tris-HCl (pH7.4), 50 mM NaCl, 1 mM DTT) or Sch9 dialysis buffer (1xPBS (pH7.4), 1 mM DTT), then concentrated with an Ultrafree-4 centrifugal filter 50K or 30K membrane (Millipore).

### *In vitro* kinase assay

To test phosphorylation of recombinant Sch9 peptides, *pho80Δ* mutant cells expressing pHA-Pho80 grown to mid-log phase, collected and lysed in lysis buffer (50 mM Tris-HCl (pH7.5), 2 mM sodium pyrophosphate, 0.1 % SDS, 1 % sodium deoxycholate, 1 % Triton X-100, 1 mM PMSF, 1 X PIC) and centrifuged at 13,000 x g for 10 min. The supernatant was incubated with anti-HA antibody and immobilized on protein A Sepharose beads (Sigma). Beads washed 3 times with lysis buffer, twice with lysis buffer containing 150 mM NaCl and then washed twice with kinase buffer (10 mM Tris-HCl (pH7.4), 10 mM MgCl2, 50 mM NaCl, 2 mM EDTA, 1 mM DTT). Beads were incubated with 20-40 μg substrate protein, 2.5 mM ATP, 0.125 μCi [γ-^32^P]ATP in kinase buffer at 30°C, 60 min. Reactions terminated with equal volumes 2X SDS sample buffer. For recombinant peptides of Whi5, *sch9Δ* mutant cells expressing pSch9-HA_5_ grown to mid-log phase, collected and lysed in lysis buffer (1xPBS (pH7.4), 2 mM sodium pyrophosphate, 0.1% SDS, 1% sodium deoxycholate, 1% Triton X-100, 1 mM PMSF, 1 X PIC, 1xPhosSTOP (Roche) and centrifuged at 13,000xg for 10 min. The supernatant was incubated with anti-HA antibody and immobilized on protein A Sepharose beads (Sigma-Aldrich). Beads washed 3 times with lysis buffer and then washed twice with kinase buffer (1xPBS (pH7.4), 4 mM MgCl_2_, 1 mM DTT). Beads were incubated with 20-40 μg substrate protein, 2.5 mM ATP, 0.125 μCi [γ-^32^P]ATP in kinase buffer at 30°C, 60 min. Reactions terminated with equal volumes 2X SDS sample buffer.

### Growth assay in hyperosmotic stress

Cells treated with 3.7 μM α-factor with either DMSO or 20 μM 1-NA-PP1 for three hours, washed with fresh media twice and then released to fresh media including 0.9 M NaCl for seven hours. Cell number assessed by OD600.

### Growth assay during recovery after nutrient starvation

Cells treated with 3.7 μM α-factor with either DMSO or 20μM 1-NA-PP1 for three hours, washed with fresh media twice and released to starvation media for three hours. Then cells were refed with normal nutrient media for seven hours. Cell number assessed by OD600.

### Plasmids

Plasmids used are in Table 2. To generate pRS413 *sch9-TEVr367-* HA_5_, pRS413 Sch9-HA_5_ was mutagenized by site-directed mutagenesis using following primers, FW-GAAAATctgtatTTTCAAGGTCCAATGATTCATAATTTAGCAC REV-ACCTTGAAAatacagATTTTCTCTAACTTGGCCTAAGAACATATG

## Acknowledgements

We thank Drs. Daniel Klionsky, Haoxing Xu, Yukiko Yamashita, Junya Hasegawa and the Weisman Lab for helpful discussions. Supported by National Institutes of Health (NIH) Grant R01-GM062261 to LSW, and Grant-in-Aid for Research Activity (KAKENHI) 21K06160 to NJ, 17H06678 to YJ, 21K06148 and 23H00382 to A. Nakano and 16H06375 to Y. Ohsumi.

## Author contributions

N. J. conceived the study; designed, performed, analyzed, and interpreted experiments; and wrote the manuscript. Y. J. designed, performed, analyzed, and interpreted experiments; and wrote the manuscript. Y. O. performed and analyzed experiments. A. Nakano and Y. Ohsumi provided critical insights. L. S. W. conceived the study; designed, analyzed, and interpreted experiments; and wrote the manuscript.

## Declared interests

The authors declare that they have no competing interests.

## Data and materials availability

All data to understand and assess the conclusions of this research are available in the main text, supplementary materials,

