## Supplemental Figure 1-3 for "A non-canonical CDK, Pho85 regulates the restart of the cell-cycle following stress"

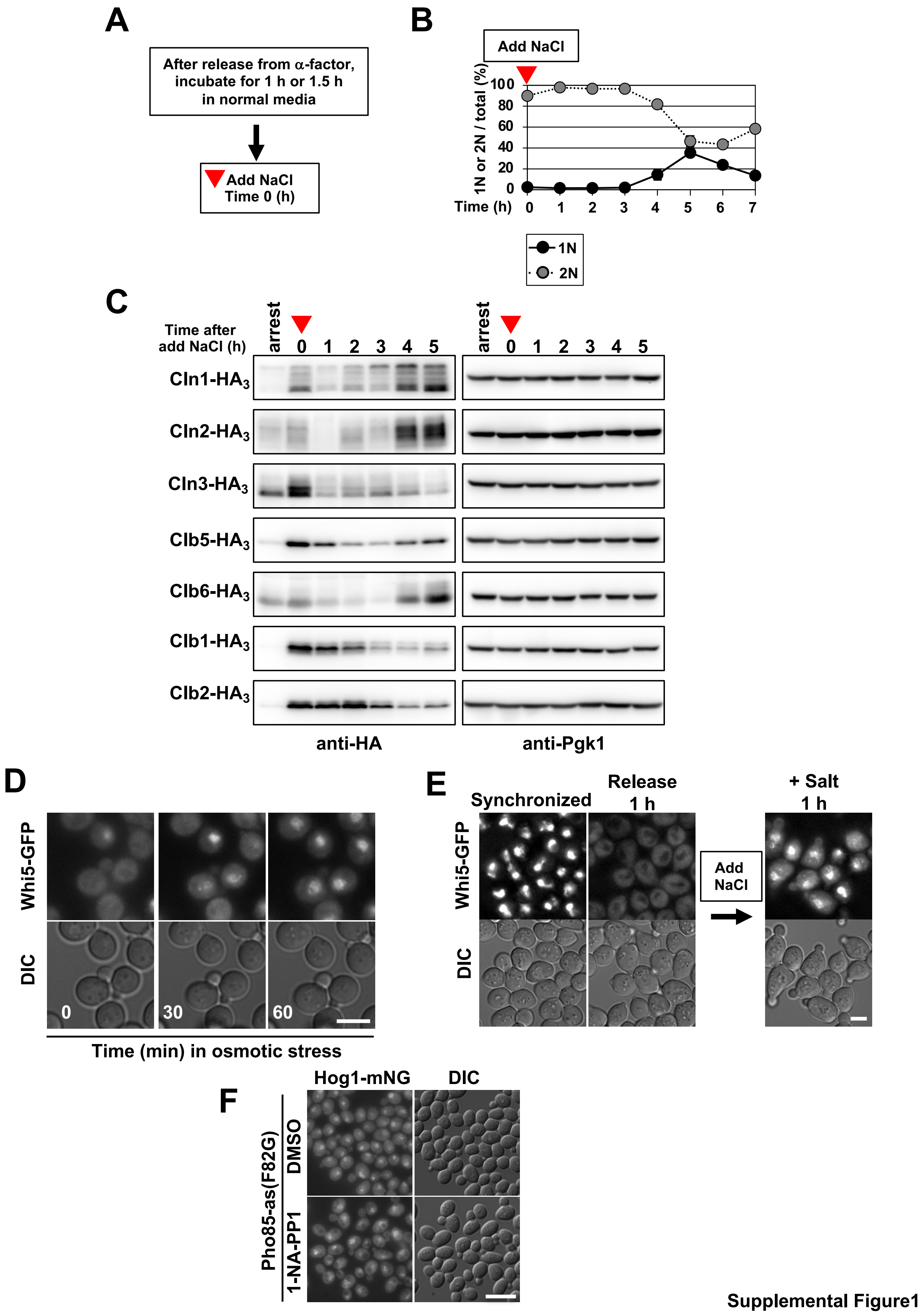


Supplemental Figure1

**Supplemental Figure 1.** (A) Schematic of experiment for (B) and (C). Indicated yeast strains arrested with α-factor with 80% of the population with 1N DNA content, were released into fresh medium for 1 h (B) or 1.5h (C) and then incubated with 0.9 M NaCl for 1-7 h. (B) DNA content was measured by flow cytometry. Mean values ± SD. (C) Western blot analysis performed using anti-HA or anti-Pgk1 (control). Blots representative of 3 independent experiments for Cln1-$HA_3$, Clb5-$HA_3$, Clb1-$HA_3$, Clb2-$HA_3$ or 2 independent experiments for Cln2-$HA_3$, Clb6-$HA_3$, Cln3-$HA_3$. (D) Following hyperosmotic stress, Whi5 is imported to the nucleus. Treatment of yeast with high salt drives Whi5-3xGFP to the nucleus at any phase of the cell-cycle. Yeast expressing Whi5-3xGFP were exposed to 0.9 M NaCl at time 0. Images were acquired every 10 min for 1 h. Bar: 5 μm. (E) Following hyperosmotic stress, Whi5 is imported to the nucleus, even in cells with buds. (F) The Pho85-as (F82G) mutant, expressing Hog1-mNG was treated with either DMSO or 20 mM 1-NA-PP1 for 3 h, then exposed to 0.9 M NaCl for 5 min. These data suggest that Pho85 activity is not required for the translocation of Hog1 to the nucleus during hyperosmotic stress. Bar: 8 μm

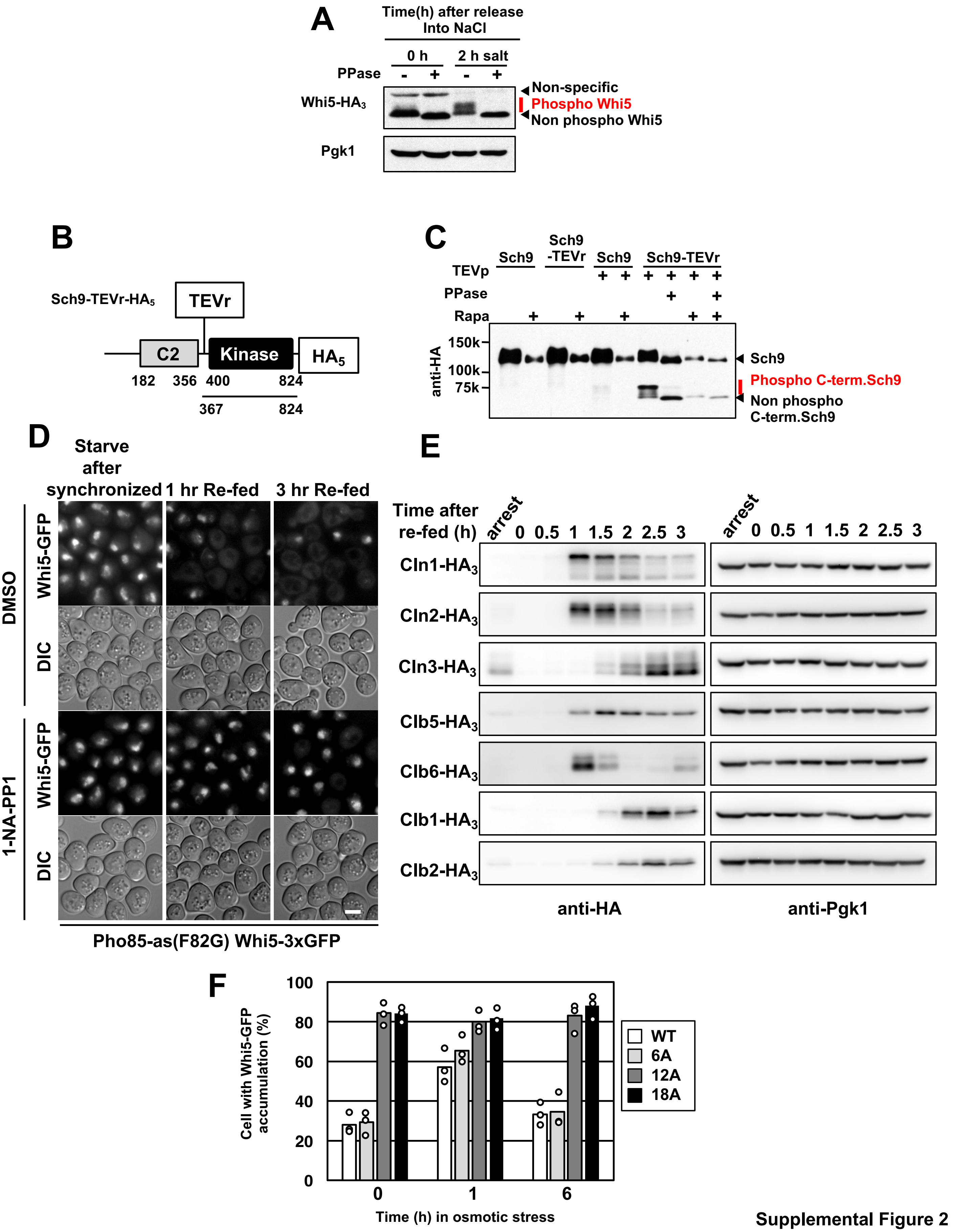


Supplemental Figure 2

**Supplemental Figure 2.** (A) When cells exit from stress-induced cell-cycle arrest at $G_1$ phase, Whi5 is phosphorylated. Yeast expressing Whi5-$HA_3$ were incubated with α-factor for 3 h. 80% of the population arrested at $G_1$ phase as assessed by DNA content. Cells were released to fresh media that included 0.9 M NaCl and then lysed with 10% TCA at the indicated times, and digested with phosphatase (PPase). Western blot: anti-HA or anti-Pgk1(control). Blots representative of 3 independent experiments. (B) Schematic of Sch9-TEVr(367)-$HA_5$. Black line (amino acids 367-824), region used in Fig. 4A and 4E. (C) The C-terminus of Sch9 is phosphorylated. Wild-type cells expressing Sch9-TEVr(367)-$HA_5$ or pSch9 treated with (+) or without (-) 0.2 µM rapamycin for 150 min, then lysed with 10% TCA, and treated with (+) or without (-) TEVp and/or PPase. Western blot analysis performed using anti-HA. Blots representative of 3 independent experiments. (D) Representative fields of cells analyzed in Fig. 5F. Bar: 5 µm. (E) Yeast expressing Cln1-$HA_3$, Cln2-$HA_3$, Cln3-$HA_3$, Clb1-$HA_3$, Clb2-$HA_3$, Clb5-$HA_3$, or Clb6-$HA_3$ were arrested with α-factor with 80% of the population with 1N DNA content, and were released into nitrogen starvation medium for 3 h and then introduced into nutrient-rich medium. Western blot: anti-HA or anti-Pgk1(control). Blots representative of 3 independent experiments. (F) The canonical-Cdc28 phospho sites are important for the export of Whi5 from the nucleus during hyperosmotic stress. As assessed in an asynchronous wild type cell expressing pRS413-Whi5-mNG, pRS413-$Whi5^{6A}$-mNG, pRS413-$Whi5^{12A}$-mNG or pRS413-$WHi5^{18A}$-mNG following hyperosmotic stress, the movement of Whi5 or $Whi5^{6A}$ to the nucleus is transient. However, $Whi5^{12A}$ or $Whi5^{18A}$ remained trapped in the nucleus. n=3 >100 cells counted per condition, per experiment.

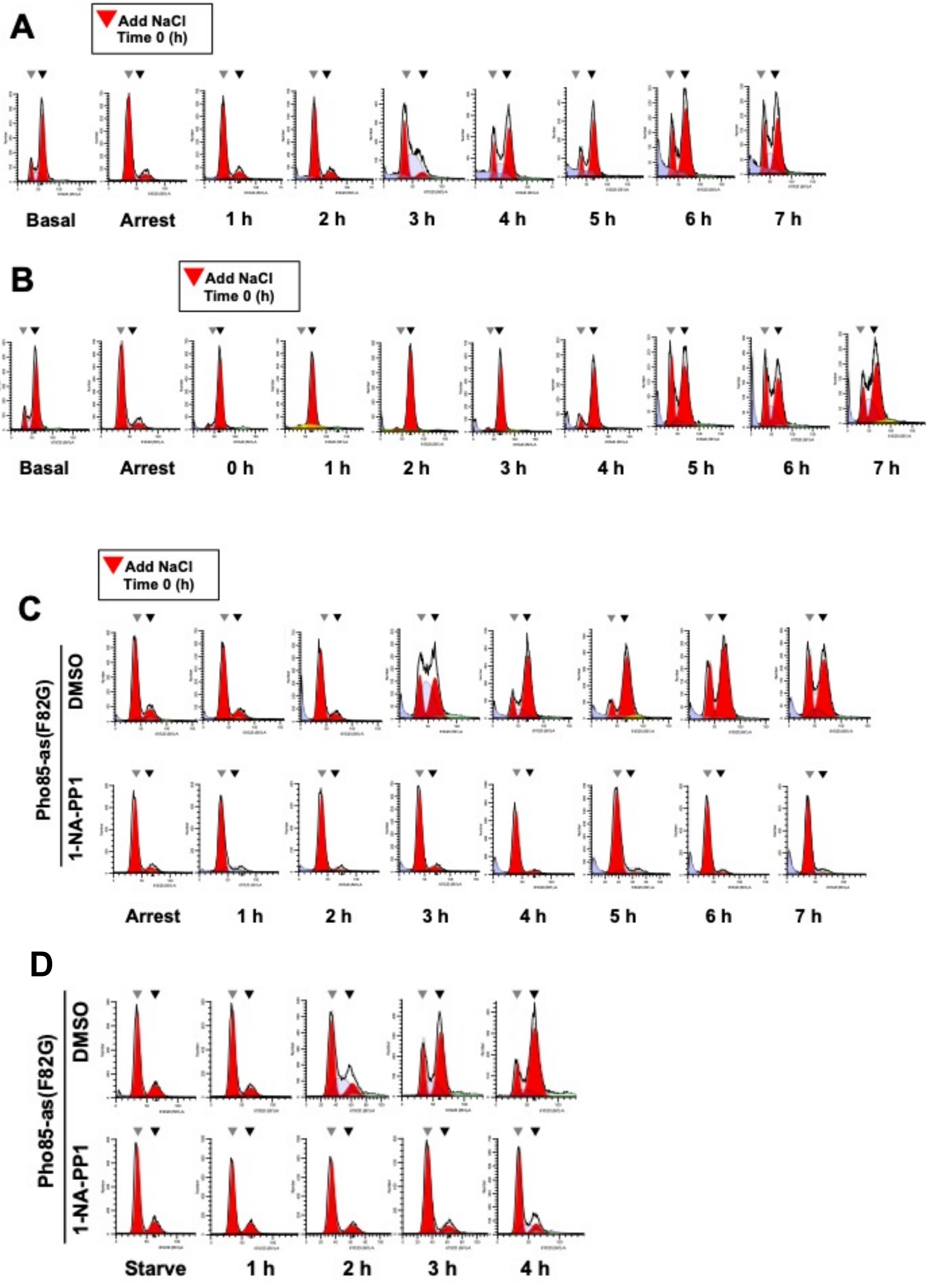


Supplemental Figure 3

**Supplemental Figure 3.** Representative histograms, (A) corresponding to Fig 1G, (B) corresponding to Supplemental Fig 1B, (C) corresponding to Fig 3A, (D) corresponding to Fig 5G. Gray triangles, peak of 1N. Black triangles, peak of 2N.
